# Tau Aggregation is Altered by Variations in its Projection Domain

**DOI:** 10.1101/2024.11.22.624898

**Authors:** Kayleigh Mason-Chalmers, Aanzan Sachdeva, Gabryelle Kolk, Justin C. Touchon, Zachary J. Donhauser

## Abstract

The intrinsically disordered microtubule-associated protein tau is known for its aberrant aggregation into neurofibrillary tangles as found in neuropathologies such as Alzheimer’s disease. This study compares three N-terminal isoforms of mutant R5L and of wild type tau to investigate how this mutation and the length of the projection domain affects aggregation behavior. Tau polymers *in vitro* were examined using atomic force microscopy imaging to compare tau filament lengths and morphologies. In a complementary analysis, the total amount of polymerization was analyzed using a Thioflavin S assay. We observed that the R5L mutation has a greater impact on filament length in shorter N-terminal isoforms of tau, whereas in longer N-terminal isoforms the mutation impacts the total amount of tau aggregation. These observations suggest that the R5L mutation affects the kinetic nucleation-elongation pathway of tau fibrillization, where the mutant impacts polymer nucleation in 2N and 1N isoforms, but has a more significant impact on elongation in the 0N isoform.

## Introduction

The microtubule-associated protein tau is primarily found in axons both in the peripheral and central nervous system, where it has important roles in cellular organization, and where it assists in the assembly, stabilization and organization of microtubules.^1–5^ Tau also has other lesser known functions such as influencing the localization of cellular kinases, interacting with the cellular membrane, and affecting the movement of motor proteins.^6–8^ Tau is widely known for its role in neurodegenerative disease,^4,5^ aggregating into paired helical filaments (PHFs) and straight filaments (SFs) which largely comprise neurofibrillary tangles (NFTs), the presence of which is associated with the class of diseases known as tauopathies. These diseases include Alzheimer’s disease (AD), frontotemporal dementia (FTD), and Pick’s disease,^2,3,9–11^ where aggregates and NFTs are highly correlated with neuronal dysfunction and death that may cause or further the progression of diseases.^1^

Structurally, tau consists of four major regions (depicted in Figure 1a): the N-terminus (or projection domain, PD), the proline-rich region (PRR), the microtubule-binding domain (MTBD), and the C-terminus.^2,7^ Additionally, tau is alternatively spliced into six major isoforms, which vary in the number of N-terminal repeats (0,1, or 2, corresponding to 0N, 1N, and 2N tau) or MTBD repeats (resulting in 3R and 4R tau).^1,5^ In the present work, the N-terminal isoforms with all four MTBD repeats were investigated. Tau isoforms with more N-terminal repeats have a higher negative charge in that region (Figure 1b), which may impact the function of the projection domain.

**Fig. 1.**
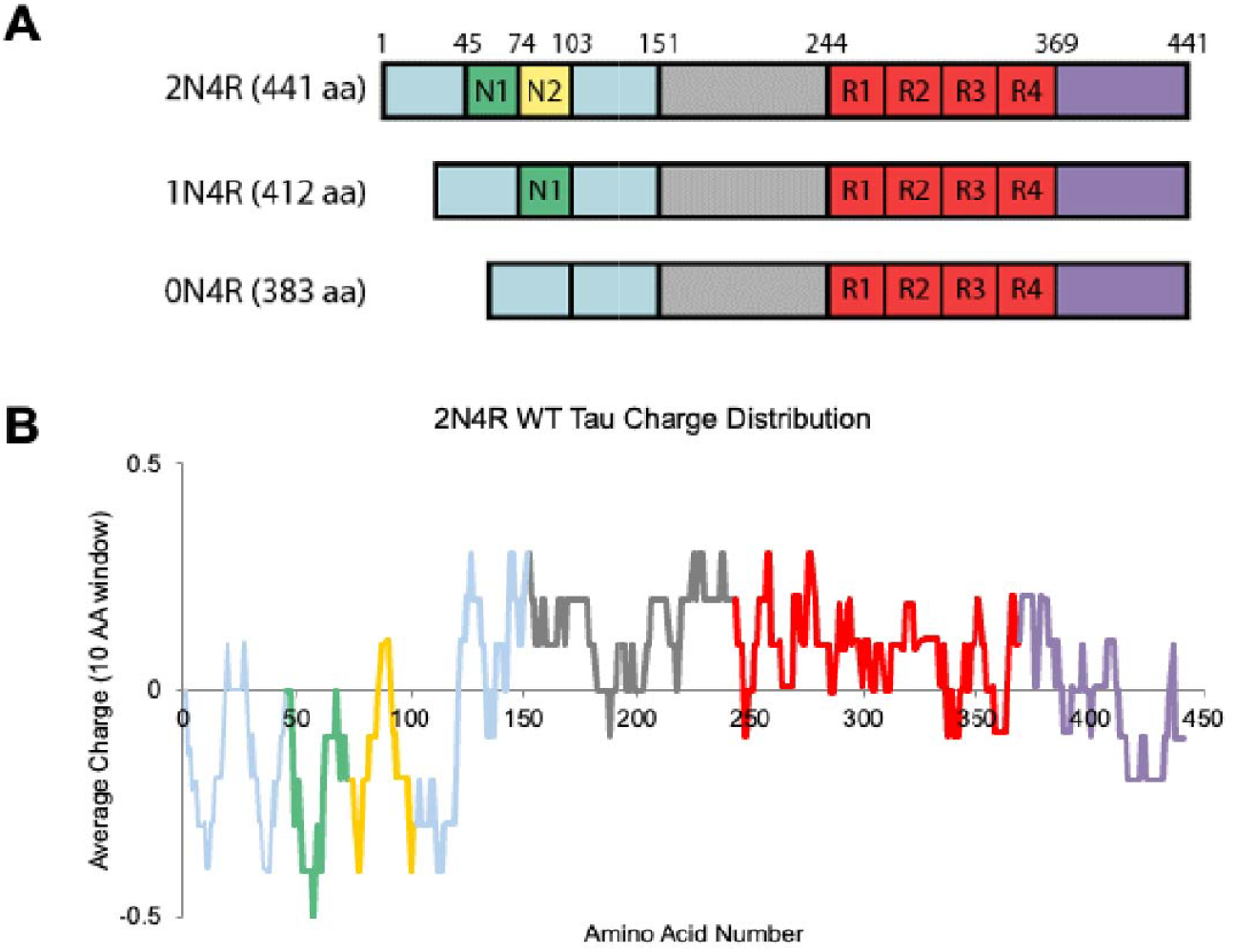
Overview of tau structure. a) Structures of 2N, 1N, and 0N4R wild type tau, which consist of the projection domain (blue, with the N1 (green) and N2 (yellow) inserts labeled), proline-rich region (grey), microtubule binding domain (red), and C-terminus (purple). Each isoform varies in its number of N-terminal repeats. b) The charge distribution of 2N4R wild type tau at pH 7.2, with charge averaged over a 10 residue window. In 1N4R the N2 (yellow) portion is eliminated, and in 0N4R both the N1 (green) and N2 (yellow) repeats are eliminated.

Physical and structural changes in the PD are key indications of dysfunctional tau,^12^ and evidence suggests that differences in the N-terminus’ length and conformation affect tau aggregation.^12,13^ The presence of the N-terminus inhibits polymerization, such that tau aggregates more readily when this region is eliminated.^14–16^ Aggregation is also influenced by the number of N-terminal repeats, as 1N isoforms tend to form more fibers than 2N,^13,17^ and 0N has been shown to form globular clusters rather than filaments.^17,18^ The lower acidity of shorter N-terminal tau isoforms are thought to increase the tendency of tau to form clusters.^18^ The focus of the present work is the R5L mutant, which decreases the acidity of this region, and is related to diseases such as familial FTD with parkinsonism in chromosome 17 (FTDP-17) and progressive supranuclear palsy (PSP).^13,17,19–21^ Other mutated tau isoforms impact aggregation of tau *in vivo*,^17^ and it is thought that R5L lowers microtubule affinity, therefore allowing aggregation through higher tau dissociation from microtubules into solution.^17,22,23^

Although tau is canonically an intrinsically disordered protein, there have been numerous studies indicating that it can adopt transient or loosely folded structures in solution, and that alterations in these structures may be related to tau pathologies. For example, Mandelkow and coworkers reported that all isoforms of tau adopt a dynamic “paperclip” conformation, where the ends of the protein fold in and the C-terminus interacts with both the MTBD and PD.^2,24–26^ Previous studies have found that R5L and MTBD mutations alter tau’s ability to form this structure,^14,27,28^ adopting more extended conformations^14^ or affecting the interaction between the MTBD, N- and C-termini.^27^ Other secondary structures of tau have been identified such as the Alz50 epitope which has interactions between the MTBD and PD,^29^ intermolecular β-sheets in the MTBD,^30^ or α-helices.^31^ When tau dissociates from microtubules, the protein fluctuates between its transient secondary structures.^14,24,32–34^ Numerous studies have suggested that tau unfolding from its secondary structures into an extended conformation is required for the formation of tau filaments.^9,14,24,32–34^

Tau forms filaments under a nucleation-elongation model (Figure 2).^9,14,24^ Monomeric extended tau first nucleates into a dimer which is initiated by the MTBD’s ^275^VQIINK^280^ and ^306^VQIVYK^311^ regions that misfold into β-strands.^25,28^ However, tau is known to misfold in numerous ways, and primary nucleation is dependent on the efficiency of tau misfolding into disease-specific dimers to form a seed,^35^ which is both rate-determining and thermodynamically unfavorable.^9,25,28,30,33^ After the seed reaches its critical size,^9,33,34^ tau monomers are added unidirectionally to form elongated straight filaments (SFs) or paired helical filaments (PHFs).^9,10,28,36–39^ The process of elongation is the thermodynamic driving force of aggregation, yet is semi-reversible, as filaments can fragment into shorter fibers but will not disaggregate into tau’s monomeric state.^40,41^ Once adequately elongated, tau fibers can cluster via higher order interactions to form neurofibrillary tangles.^9^

**Fig. 2.**
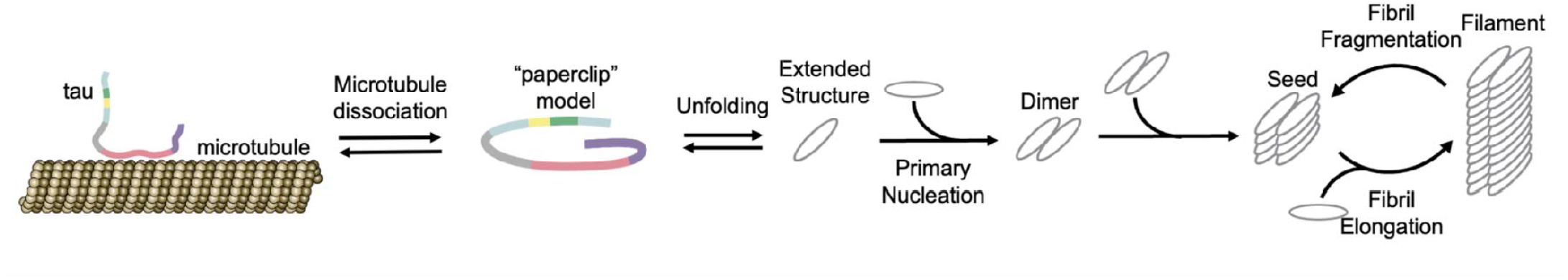
Nucleation-elongation pathway of tau into fibers. When tau dissociates from the microtubule and forms one of its secondary structures (paperclip shown in this figure), it can either reassociate or conformationally change in order to nucleate and then elongate into paired helical filaments or straight filaments.^9,19,41^

Tau filaments generally consist of a stable core comprised by hydrogen-bonded β-strands of the R3 and R4 repeats.^34,35,42–44^ These fibers vary in cross-sectional core conformations, and are unique for different tauopathies,^35,45–53^ leading to a general hypothesis that these distinct folding tendencies lead to unique tauopathies.^16,18,35,39^ Most structural studies lack information on the N-terminus,^45–52^ as the PD largely participates in the disordered “fuzzy coat” that surrounds the core of fibrils,^34,35,42–44^ and eludes characterization from techniques such as cryo-EM. There is some evidence suggesting an interaction of ^7^EFE^9^ of the N-terminus with L317 and L321 of the core,^34,35^ which is relevant when considering the R5L mutation’s close proximity to the ^7^EFE^9^ motif which may impact the formation or stability of fibers.^35^

Given that PD mutants such as R5L are found in specific diseases,^54^ it is plausible that this mutation causes tau fibers to fold uniquely compared to WT.^51,55,56^ Compared to previous studies that use electron microscopy with longer isoforms of R5L and WT tau, we conduct a complementary analysis with atomic force microscopy (AFM), examining the three 4R N-terminal isoforms of tau. In contrast to previous studies, we focus our analysis on fibers polymerized for a longer time period (four days), providing new insights into any differences between “mature” fibers across the variants studied. We aim to deepen our comprehension of how R5L acts on a monomer, fibril, and macroscopic level. Our results show that the projection domain affects tau folding, implicating its role in the nucleation-elongation model. A better understanding of the role of the projection domain in aggregation and in filaments’ fuzzy coat may elucidate of how various projection domain mutants and isoforms affect the progression of these diseases.^55^

## Materials and methods

### Cloning and tau purification

2N4R tau (UniProt ID P10636) was obtained as previously described.^2^ Mutagenesis to produce additional isoforms and mutants was performed using the Q5 Site Directed Mutagenesis Kit (New England Biolabs). 1N4R tau was generated using by an initial round of mutagenesis with primers (5’- CTGAAGAAGCAGGCATTGG-3’ and 5’-CTTCCGCTGTTGGAGTGC-3’) designed to eliminate one N-terminal insert (residues 74-103 in 2N4R tau), and 0N4R tau was generated with primers (5’- CTGAAGAAGCAGGCATTGGAG-3’ and 5’-CTTTCAGGCCAGCGTCCG-3’) that eliminated both N-terminal inserts (residues 45-103). The R5L mutation was added to each WT isoform using primers (5’- ATGTGACAGCACCCTTAGTG-3’ and R: 5’-CTTTCAGGCCAGCGTCCG-3’). Template digestion and PCR product ligation were completed using the KLD Enzyme Mix standard protocol (New England Biolabs). The PCR products were transformed into DH5α competent cells and incubated overnight on LB-agar plates containing 100μM ampicillin. For each tau isoform, several single colonies were selected and plasmids purified using a QIAprep Spin Miniprep Kit (QIAGEN). DNA sequences were confirmed with sequencing from Molecular Cloning Labs.

The mutated plasmids were transformed into One Shot BL21(DE3) Chemically Competent *E. coli* cells (Thermo Fisher) and incubated overnight at 37°C on LB-ampicillin plates. Tau purification was conducted as previously described,^2^ with cells lysed using sonication on ice rather than French Press. Following the purification, tau concentrations were typically 0.1-0.5 mg/ml in BRB8 (8 mM K-Pipes, 0.1 mM EGTA, 0.1 mM MgCl_2_, pH 6.8), and were lyophilized and stored at -20 °C until use. Following purification, protein concentration was determined by micro-BCA assay with BSA as a standard.

### Tau aggregation

Arachidonic acid (ARA, Nuchek Prep) was aliquoted in ethanol and the solvent was blown off using nitrogen gas. Aliquots were stored for up to 3 months at -80 °C, and immediately prior to use, they were resolubilized in ethanol at 10 mM. Lyophilized aliquots of tau were rehydrated with water to 1/10 of their original volume and then diluted to a final concentration at 5 μM or 10 μM in polymerization buffer (10 mM HEPES, 100 mM NaCl, 0.1 mM EDTA, 5 mM DTT, pH 7.64), with ARA in a 37.5:1 ratio to tau.^13^ Polymers were allowed to grow for four days in darkness at room temperature.

### AFM in-air imaging

30μL of polymerized tau sample was deposited onto freshly-cleaved muscovite mica (V-1 quality, Electron Microscopy Sciences) and adsorbed for 3 minutes. The sample was then rinsed with ∼3mL ddH_2_O and dried using nitrogen gas. AFM imaging was conducted with a Bruker MultiMode AFM operating in intermittent contact mode, with POINTPROBE Silicon SPM Sensor cantilevers (42 N/m, NanoWorld).

Tau fibers for each image were individually selected, traced, and measured using a custom MATLAB script. The area covered by tau aggregates was determined after flattening and thresholding the image using a custom MATLAB script. We used a two-way ANOVA to analyze the effects of tau mutants and isoforms on fiber length. Fiber length data were log-transformed to normalize the data distributions. A post-hoc test was used to compare the effects of mutant vs WT on fiber length separately for each variant, with data analysis conducted in R v4.3.3.^57^

### Fluorescence assay

After four days of polymerizing undisturbed in the dark, 1mM Thioflavin S (ThS, Sigma Aldrich) was added to each solution and measured using a Synergy HTX Multi-mode Microplate Reader (Biotek) using a 485/20nm excitation filter and 528/20nm emission filter.

## Results

### R5L vs WT tau dry imaging

To define the terminology used in this paper, we use “fibers” or “fibrils” to describe isolated, well-structured polymers as observed in AFM images, “clusters” are groups of fibers that have accumulated to become indistinguishable with some fibril ends visible around the edges, and “aggregates” is inclusive of both fibers and clusters. AFM images were collected after four days of polymerization, which contrasts with other studies that only reported data on R5L up to only 24 hours.^9,17,19,28^ The density of protein observed in AFM images depends on the species’ ability to bind uniformly to a well-defined surface (in this case mica), and the relative sample size is very small in comparison to the overall population, which results in some variability in the AFM images. To mitigate this, multiple images were taken of each sample in order to find representative areas of the surface.

To optimize AFM images for our analyses, we tested a range of solution conditions, and found that deposition from 5 μM for 3 minutes yielded surfaces that typically contained numerous, yet well-dispersed, fibers for all variants. Several of the variants were sampled and imaged multiple times, and with each showing qualitatively similar results across samples. For the analyses presented here, three images were collected from a single, protein sample to maintain consistency for each variant.

Representative images are depicted in Figure 3, which reveal the total amount of fibers on the surface and their tendency to cluster. The clustering phenomenon is mostly seen in 0N for both wild type and the mutant (clusters indicated by red arrows in Figure 3), although 1N was observed to cluster in some images.

**Fig. 3.**
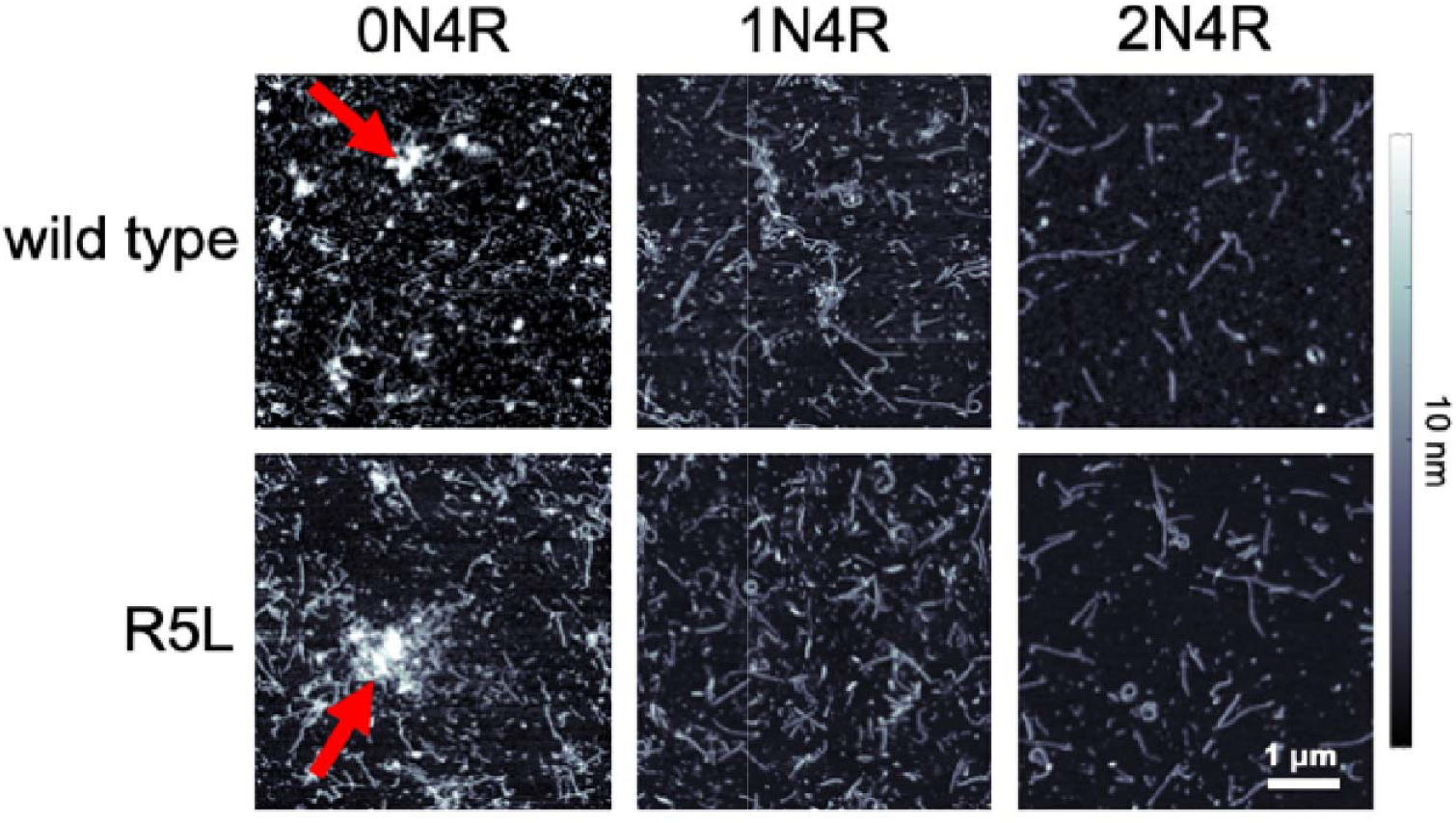
AFM images of tau fibers. Polymerization solutions containing 5μM tau were incubated for four days and then deposited on mica for AFM analysis. The red arrows indicate two examples of tau clusters.

The length analyses for each protein are shown in Figure 4, which showed a trend of longer isoforms of tau forming longer fibers. Fiber length was significantly affected by mutant and isoform (two-way ANOVA, mutant: *F*_1, 1796_ = 22.0, *p* < 0.001, variant: *F*_2, 1796_ = 140.2, *p* < 0.001). There was also a significant interaction between mutant and isoform, such that differences between mutants were dependent on isoform (two-way ANOVA, *F*_2, 1796_ = 3.2, *p* = 0.04). When comparing mutant vs. WT, the mutation only significantly affected the average length of fibers for 0N4R tau (*p* < 0.001) and not 1N (*p* = 0.07) or 2N4R (*p* = 0.70).

**Fig. 4.**
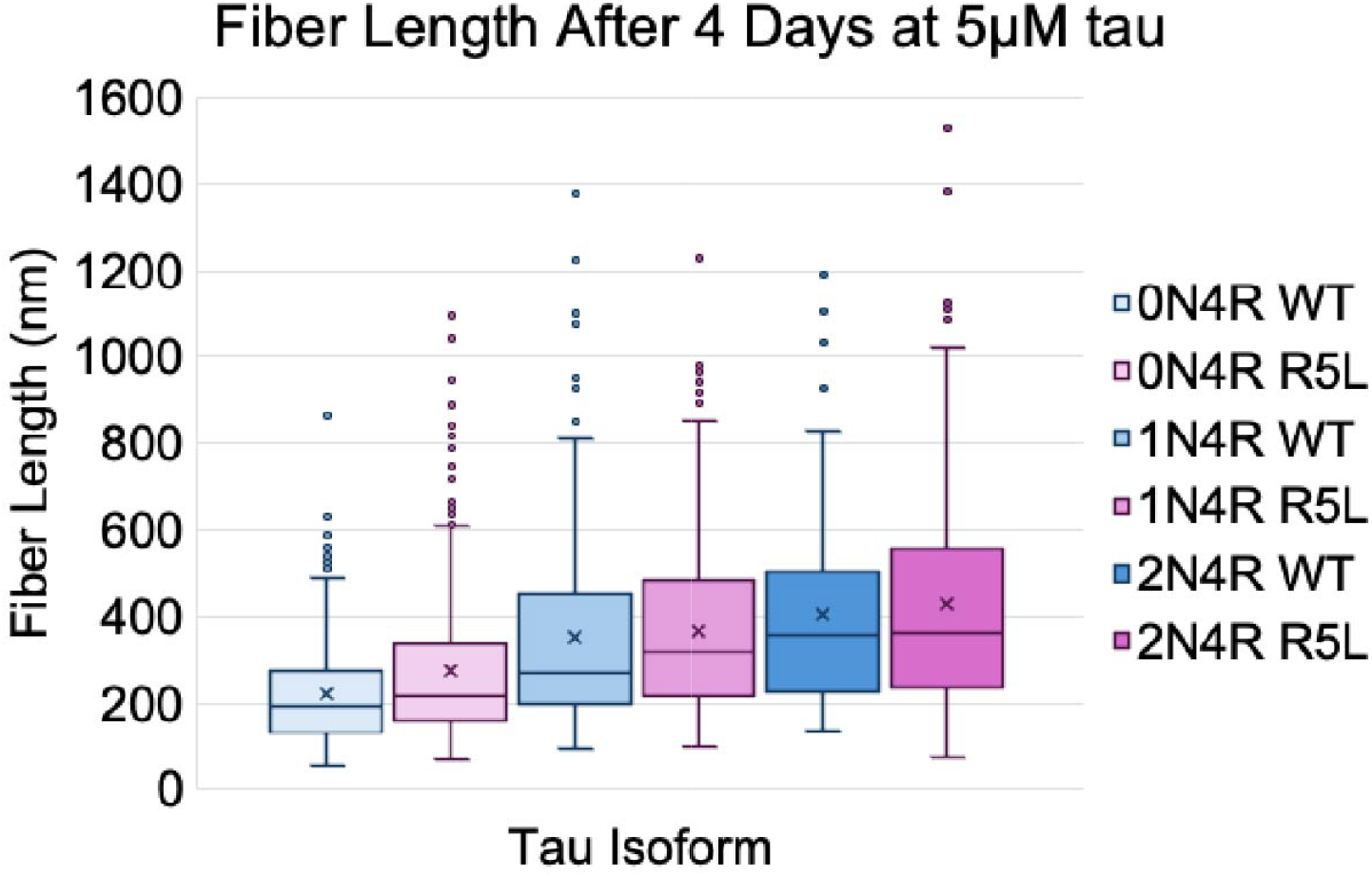
Fiber length distribution of tau variants. The fiber lengths were gathered by manually selecting distinguishable fibers on three different 5μm AFM images. X represents the sample mean.

The amount of tau aggregation in the AFM images was assessed using two separate methods. First, the total sum of the lengths for each image was measured and averaged (Figure 5a). However, in images with many larger aggregates and clusters, it was difficult/impossible to distinguish and accurately measure each individual fiber, thus underestimating the total aggregate density in those images.

**Fig. 5.**
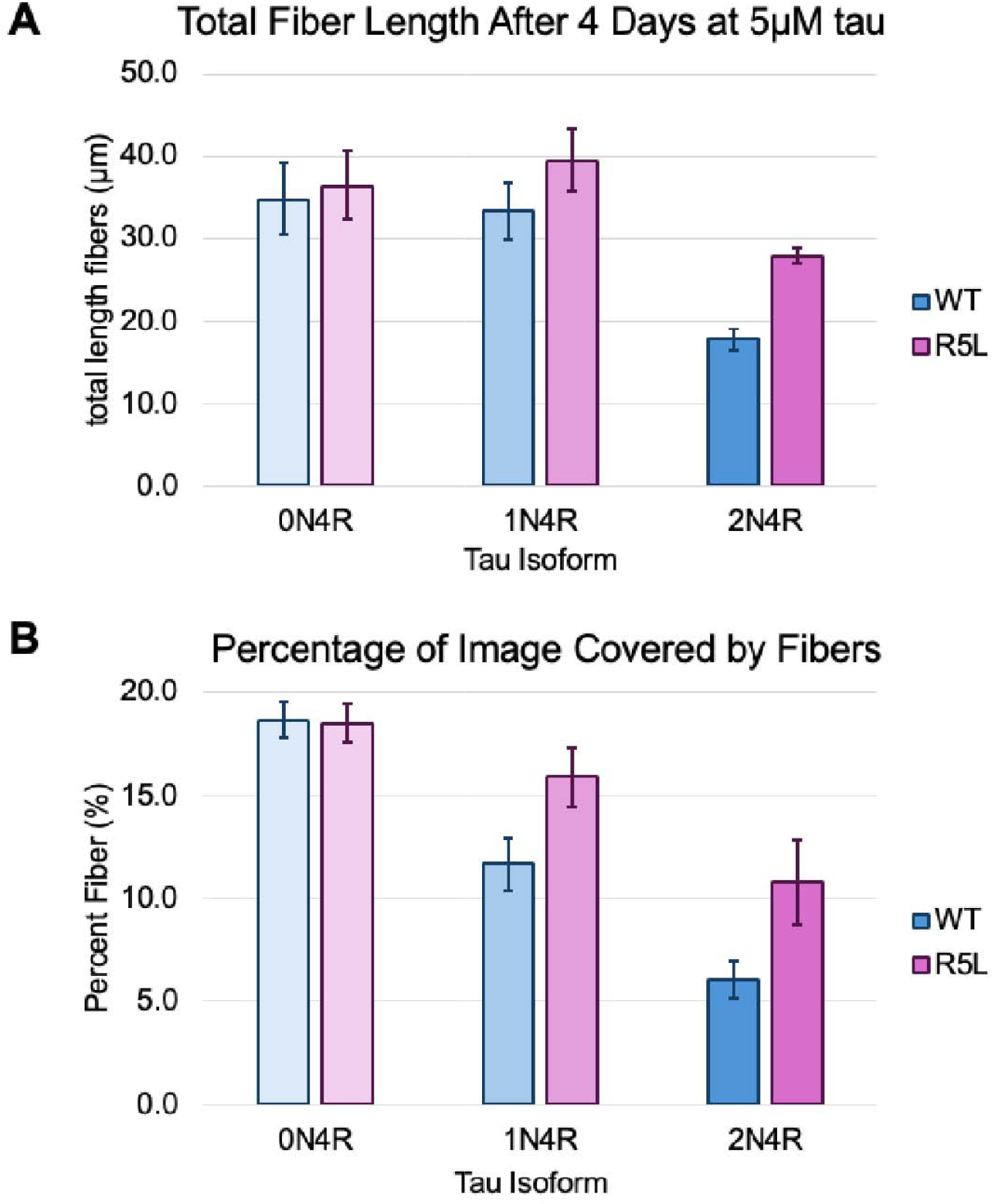
Analysis of total polymer in AFM images. a) The total length of tau fibers for each variant in 5μm images, averaged over 3 images, along with standard deviations. b) The percentage of the image covered by tau aggregates for each variant. The total area covered by tau aggregates was determined for each 5μm image and normalized to the total area of the image, and averaged over 3 images.

Therefore, as a second estimate of total tau polymerization, the total area of the surface covered by protein was measured and taken as a percentage of the image (Figure 5b).

When only taking into consideration the distinguishable fibers (i.e. the total length measurement, Figure 5a), 0N4R and 1N4R had similar amounts of fiber, with 2N4R having less total fiber length. Whereas, when considering total surface coverage, 0N4R aggregates more than 1N4R (Figure 5b). These data indicate that there is the same amount of distinguishable fibers for 0N4R and 1N4R, where the extra area accumulated by 0N4R is clustered protein and is unassignable as individual fibers in our data. Consistent with the data in Figure 5, a previous report showed that in WT tau, 1N4R aggregates around 50% more than 2N4R, and 0N4R tau has the highest total aggregation.^18^

Based on our data, the mutation of each isoform had an opposite influence on total aggregation (Figures 5a and b) as compared to fiber length (Figure 4). Whereas the average length of 0N R5L fibers was significantly longer than 0N WT (Figure 4), we observed that they form about the same total amount of aggregate (Figure 5a and b). On the other hand, R5L mutants for both 1N and 2N tau form similar length fibers to WT (Figure 4), yet for both isoforms R5L tau forms significantly more aggregates (Figure 5a and b). In summary, we observed that the R5L mutation has a greater effect on the lengths of individual fibers of the shortest isoform, while the mutation produces more total aggregation for 1N and 2N4R and not 0N4R.

### Results of fluorescence assay

Total tau aggregation was also measured in triplicate using ThS fluorimetry after four days of growth (Figure 6). This assay showed similar results to AFM imaging when comparing mutants of the same isoform (Figure 5b). The fluorimetry results for 2N4R R5L are consistent with the trends observed in our AFM data, and with previous fluorimetry and LLS assays reporting that R5L increases the total aggregation for 2N tau.^19,28^ For both the 1N and 0N isoforms, our fluorescence data indicate that R5L may aggregate slightly more that WT, which is consistent with our AFM total aggregation data, but differs from previously published data^17^ on these shorter isoforms.

**Figure 6.**
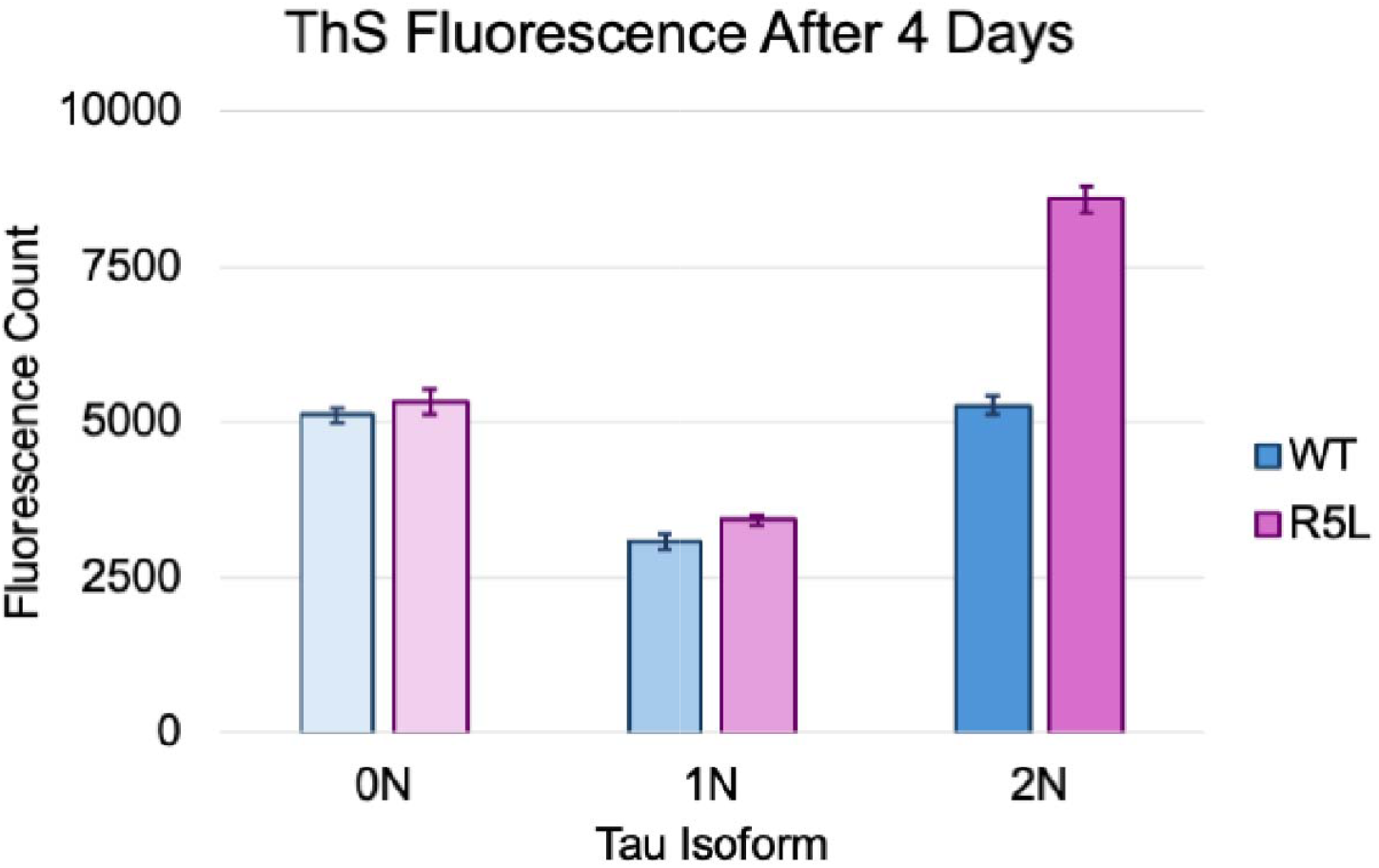
Total ThS fluorescence. Fluorescence was measured for each tau variant after 4 days at 5μM. Each bar represents an average of three measurements.

When comparing total fluorescence across isoforms, it is notable in our data that 1N R5L and WT produced the lowest total signal, while there was more total apparent aggregation from both 0N and 2N. These data contrast with previous reports that had observed a much smaller difference in total fluorescence across isoforms.^17^ The trends in our fluorescence data across isoforms also differs slightly from our AFM data, where we observed that longer isoforms produced somewhat less total aggregation.

## Discussion

### Comparison of AFM and fluorometric results

Multiple previous studies of tau aggregation have found disparities between electron microscopy images and ThS fluorescence for the N-terminal isoforms of tau. These discrepancies between imaging data and fluorimetric data may be explained by contrasting the limitations and strengths of the two approaches. As previously mentioned, AFM imaging is limited by the ability of the protein and fibers to bind to the mica surface as well as the sample size being very small compared to the total population of fibers. In comparison, fluorescence may be able to detect very small fibers that are ThS-sensitive yet not visible at the resolution of our AFM images. The main caveat of the fluorimetry assay is that it depends on the interactions of ThS with filaments. ThS binds to PHFs more readily than SFs, and so it may be more sensitive to variants that favor PHFs over SFs,^58^ conformations which we are unable to distinguish in our AFM imaging data, though this could be possible using higher resolution AFM or SEM.^18,59^

Although R5L is a relatively small mutation, it is possible that ThS binding differs between the mutant and wild type tau, or between tau isoforms. In addition, ThS binds more readily to the middle portions of fibers than the ends, so it is more effective in measuring longer fibers.^60^ Additionally, ThS may be excluded from the inner portions of large clusters of fibers as found for 0N and 1N tau, thus effectively reducing the relative measured fluorescence in those samples. Taken together, these effects may explain why 2N tau has the highest fluorescence as compared to shorter isoforms, as 2N forms the longest fibers and fewest clusters (Figures 3 and 6). In the future, a comparative study using other dyes, such as Thioflavin T or congo red, could further inform this study as they may bind more effectively to all tau fibrils formed, and may be a more consistent reporters of tau polymerization.^9,58,61^

### The effect of isoform on tau aggregation

Previous studies of R5L and WT tau isoforms have produced a varied picture of both isoform and mutant polymerization, and so it is important to place this study within the context of these reports. Some important factors to consider when comparing to earlier studies include the concentrations and ratios of tau to ARA, as well as the incubation times used for polymerization.

As previously reported, 0N tau has been found to undergo its nucleation step and form few elongated fibers, resulting in shorter filaments, and more globular or afilamentous aggregates.^13,17,18^ We have, on some occasions, observed that purification of 0N tau produces protein that does not polymerize into obvious filaments when induced with ARA, in spite of similar purity (assessed by SDS-PAGE), storage conditions, and age when compared to successful purifications (Supporting Information Figure 3). Yet, the fibers observed in Figure 3 for 0N4R WT and R5L contrasts previous reports that 0N tau will not fibrillize with ARA as an inducer.^17,18^ As the presence of the N-terminal is inhibitory to aggregation,^14–16^ a simple model would expect shorter N-terminal tau isoforms should be able to aggregate more readily.

This idea is consistent with our total aggregation data (Figure 5b), and it is further evident in our AFM data (Figure 3 and Supporting Information Figure 3) that 0N inherently produces a variety of aggregate types, both fibers and globular structures. This presence of larger clusters may, in part, be caused by the higher number of individual fibers formed with 0N tau, and indeed at higher concentrations we observed more clustering (Supporting Information Figure 3). Additionally, a higher propensity for clustering may be an inherent characteristic of 0N tau.

### The effect of the R5L mutant on tau aggregation

Previous EM studies have also had varying results of fiber length and total aggregation for R5L versus wild type tau.^9,17,19,28^ Of these studies, three only consider 2N4R tau. One study found that 2N4R R5L tau forms longer fibers with the same amount of total aggregate after incubating overnight,^29^ however, the same lab also reported that 2N4R R5L has the same length and total aggregation as wild type under the same conditions.^23^ The conditions under which tau is polymerized also affects the characteristics of the resultant fibers, and indeed Gamblin *et al*. also found that 2N4R R5L forms shorter fibers and more total aggregate compared to WT when using half the ratio of arachidonic acid:tau and a polymerization time of five hours.^17^ Using the aggregation inducer thiazine red rather than ARA, Chang *et al*. found that after 24 hours all concentrations of tau ranging from 0.2 μM to 0.8 μM show 2N4R R5L forms the same length fibers as wild type, however R5L forms more total aggregate.^9^ The results reported herein are most consistent with the study by Chang *et al*.^9^

A previous study using all three N-terminal isoforms of 4R tau similarly agrees with part of the conclusions from this experiment.^17^ Although this previous work found that 0N tau did not form elongated fibers, they did find that 0N R5L formed fewer total aggregates than WT and fibers were shorter in length. For the 1N isoform, the previous study found that the R5L mutation did not affect either the total aggregation or length of filaments,^23^ whereas our work observes the mutation increasing the aggregate mass. We note that there are differences in the conditions used previously and those used here, as we used 5 μM tau concentration and a 5 day incubation, while Mutreja *et al*. used a concentration of 2 μM and overnight incubation.^23^

### The role of the projection domain in tau’s nucleation-elongation pathway

Based on our combined AFM and fluorimetry results, the R5L mutation increases the length of 0N4R fibers while not affecting its total amount of aggregation. In contrast, for the 1N and 2N isoforms, the R5L mutation does not significantly alter fiber length yet increases the total amount of aggregation. Since R5L forms more fibers for both 1N and 2N, but of the same length as WT, we hypothesize that the mutation primarily affects the nucleation step of tau’s aggregation pathway for longer isoforms. Similarly, only the fibril length of 0N tau is significantly affected by the R5L mutation while forming the same total amount of fibrils, suggesting that the mutation alters the elongation process of fibrillization for the shorter 0N isoform. Therefore, the data presented herein may indicate that the R5L mutation more strongly impacts the nucleation of longer isoforms and the elongation of shorter isoforms.

Based on the data presented here, the N-terminal deletions (between 2N, 1N and 0N) likely play the biggest role in altering the nucleation steps of tau, with the R5L mutation having a smaller impact on nucleation. Because the N-terminus inhibits aggregation,^32,62^ it is not surprising that the shorter isoforms could permit a greater total amount of aggregation.^16^ We hypothesize that the decrease in negative charge in the projection domain region as found in 0N4R and 1N4R tau may destabilize polymerization-resistant secondary structures such as the paperclip conformation, therefore leading to more nucleation followed by aggregation.^9^ When comparing WT to R5L for 0N4R and 1N4R, they are differentiated primarily on the basis of isoform (and less by mutation), such that within each isoform there does not appear to be significant mutant-WT differences in total aggregation in our long-term AFM and fluorimetry data.

Whereas for the 2N4R isoform, the R5L mutation may have a greater destabilizing effect on the paperclip conformation and/or other transient secondary structures.^32^ Destabilization of polymerization-resistant conformations by the R5L mutation^28^ would bias the protein towards unfolding and nucleation, which would result more total aggregation as observed here and by others.^9,30,33^ This is consistent with earlier work by Chang *et al*., who found that 2N4R R5L has an increased nucleation rate but no change in elongation.^9^

While the primary focus of this work was on the examination of mature fibers after several days of growth, we did also study the initial stages of fiber growth using a fluorometric kinetic assay over the initial four hours of polymerization. We parameterized the 4-hour fluorimetry data with a modified version of the Avrami equation to find the rate constants of the initial growth. In the presence of ThS, the initial polymerization of R5L was faster across all isoform pairs (Supporting Information Figure 4 and Table 1). Under our conditions, in the presence of both ARA and tau, ThS signal increased immediately and rapidly upon addition of inducer, with no observable lag time, while control experiments lacking either ARA or tau showed no increase in fluorescence. Further, fitting the data revealed Avrami exponents around 0.5, which is consistent with diffusion-limited growth of 1-dimensional fibers^63–65^ that would be expected for rapid fiber growth. Our data is unlike previous reports using fluorimetry, where a measurable lag time is consistent with slow initial nucleation. The rapid rise in fluorescence, combined with the presence of tau seeds lacking long fibers in 4-hour AFM images (not shown), lead us to conclude that, in addition to fiber growth, ThS is sensitive to other ARA-tau interactions that occur immediately and progress rapidly.

Isoforms with longer projection domains form longer fibers, and we hypothesize that there are two complementary effects that contribute to this behavior. Previous work with cryo-EM found that there are stabilizing interactions between the negatively-charged projection domain and the positively-charged filament core, with the ^7^EFE^9^ motif participating in a key interaction.^35^ Thus, the higher negative charge density of the projection domain in longer isoforms may result in more favorable electrostatic interactions with the filament core, resulting in the formation of longer fibers. For a similar reason, we posit that the R5L mutation, in close proximity to the ^7^EFE^9^ motif,^35,54^ would also have a stabilizing effect that favors longer fibers. In our data, the specific stabilization caused by the R5L mutation does not lead to statistically significant changes in fiber length for the 1N or 2N isoforms, but does so for the 0N isoform. This may be due to the greater relative effect of a single mutation on the acidity of this region in the shorter projection domain.

Future studies, including multiple days of AFM imaging combined with longer term time-based fluorimetry measurements could yield additional insights into the kinetics of tau aggregation. As previously mentioned, the smaller fluorescent dye Thioflavin T binds more readily to tau fibers,^58^ and can be used to study kinetics.^30,55,61^ This work used 5μM tau concentration during polymerization, which is similar to intracellular tau concentrations of 2-4μM,^13^ however using smaller concentrations of tau can help to better compare to previous reports.^9,17,18^ To form filaments closer to those found in disease, recent work has reported variations in tau fiber structures using different polyanion inducers.^55^ By using these techniques to form aggregates similar to those collected *in vivo*, the information gathered by *in vitro* experiments may be more disease-relevant. Similarly, to differentiate these filaments, conducting AFM imaging in liquid environments may help better understand the hydrated fiber structure, how tau fibers interact, and whether PHFs or SFs are formed by specific tau variants.^32,59^

## Conclusions

The projection domain was observed to significantly impact the formation of tau aggregates. Shorter isoforms of tau formed the greatest number of aggregates, yet longer isoforms formed longer fibers. Using the R5L mutation as a lens to study the PD, we hypothesize that the mutation affected the nucleation of longer isoforms due to its impact on tau’s transient structure in solution and elongation of shorter isoforms by impacting the interactions between the PD and the filament core.

Based on previous results and those presented in this study,^13,17,18,28^ it is clear that *in vitro* tau aggregation studies are very sensitive to experimental parameters, including concentration of tau and ARA, the type of measurement used, and incubation time. As the results of inducer-based tau aggregation are quite sensitive to experimental conditions, there is an opportunity to use different inducers and conditions to probe mechanisms of tau aggregation, however, it also complicates the interpretation of the physiological significance of these types of aggregation studies. Future studies exploring the exact mechanism of these inducers with tau may elucidate why small differences have an outsized impact on experimental results.

A better understanding of the pathways of aggregate formation is essential for future inhibitive drug designs.^66^ For example, they could focus on tau nucleates, or seeds, that are known to be able to spread throughout the brain and infect other regions of the brain.^66^ This study provides evidence that shorter projection domain isoforms have a greater aggregation propensity and form the most amount of seeds, though aggregation propensity does not seem to correlate with higher seed spreading.^67^ Previous studies have shown various diseases having different ratios of 3R:4R tau,^4,5^ and perhaps ratios of 0N:1N:2N tau are also important. Further study into other PD mutants such as R5H may further elucidate the importance of the projection domain in aggregation as found in this study.^21^

## Supporting information

all supplemental figures

## Acknowledgements

This work was made possible by the NIH grant #1R15N5108245-01A1 and the Vassar College Undergraduate Research Summer Institute (URSI) program. Thank you to Shira Freilich, Shaharia Khan, Micaela Primer, and Lukas Johnson for providing some of the purified proteins used in this study.

## Supporting Information

**S1 Fig. Representative fiber length analysis images**. Fiber length analysis of the representative images from Figure 6 using a custom MATLAB script. Each indistinguishable full fiber was manually traced and selected.

**S2 Fig. Representative area analysis**. Area analysis of the representative images from Figure 6 using a custom MATLAB script. The aggregate area is highlighted in red and was used to find the percentage of the image covered by fibers.

**S3 Fig. Variable results from 0N4R polymerizations**. 0N4R R5L fibers at two different concentrations from two different purifications that yielded fibrillization and aggregation without fibrillization. All images are after four days of growth at 5 μM and 10 μM.

**S4 Fig. Initial 4-Hour ThS Fluorescence of Tau Variants**. a) 0N4R b) 1N4R and c) 2N4R R5L (pink) and WT (blue) fluorescence signal for the first four hours of polymerization in the presence of Thioflavin S. Points represent an average of three replicates, solid lines represent fits to the Avrami equation (see details below) and shading represents one standard deviation.

Polymerization solutions without tau (Polymerization Buffer, ARA, and 1mM Thioflavin S) were added in triplicate to wells in in a clear-bottom black 96-well plate (--). The Synergy HTX Multi-mode Microplate Reader (Biotek) was prepared for use and ready before adding tau (5μM) to the prepared wells using a multichannel pipette. The plate was immediately inserted in the plate reader and run for four hours with bottom fluorescence being measured at excitation 485/20 emission 590/35 every three minutes. Using this method, the time between adding tau and the first reads was on the order of 30-90s.

**S1 Table. Summary of fitting parameters**.

Time-based fluorometry data was fit to a modified version of the Avrami equation:

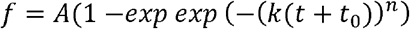

Where *f* is the normalized fluorescence and *t* is time. The coefficient *k* is a rate constant dependent on both the nucleation and growth rates, and *n* depends on the dimensionality of the polymerization (*n* = 0.5 is expected for instantaneous nucleation and diffusion limited growth in one dimension).^68^ *t*_*0*_ is included to account for the time delay between mixing solutions and the first recorded measurement.

