## Supplementary figures and images for "Tau Aggregation is Altered by Variations in its Projection Domain"

### all supplemental figures

#### Slide 1
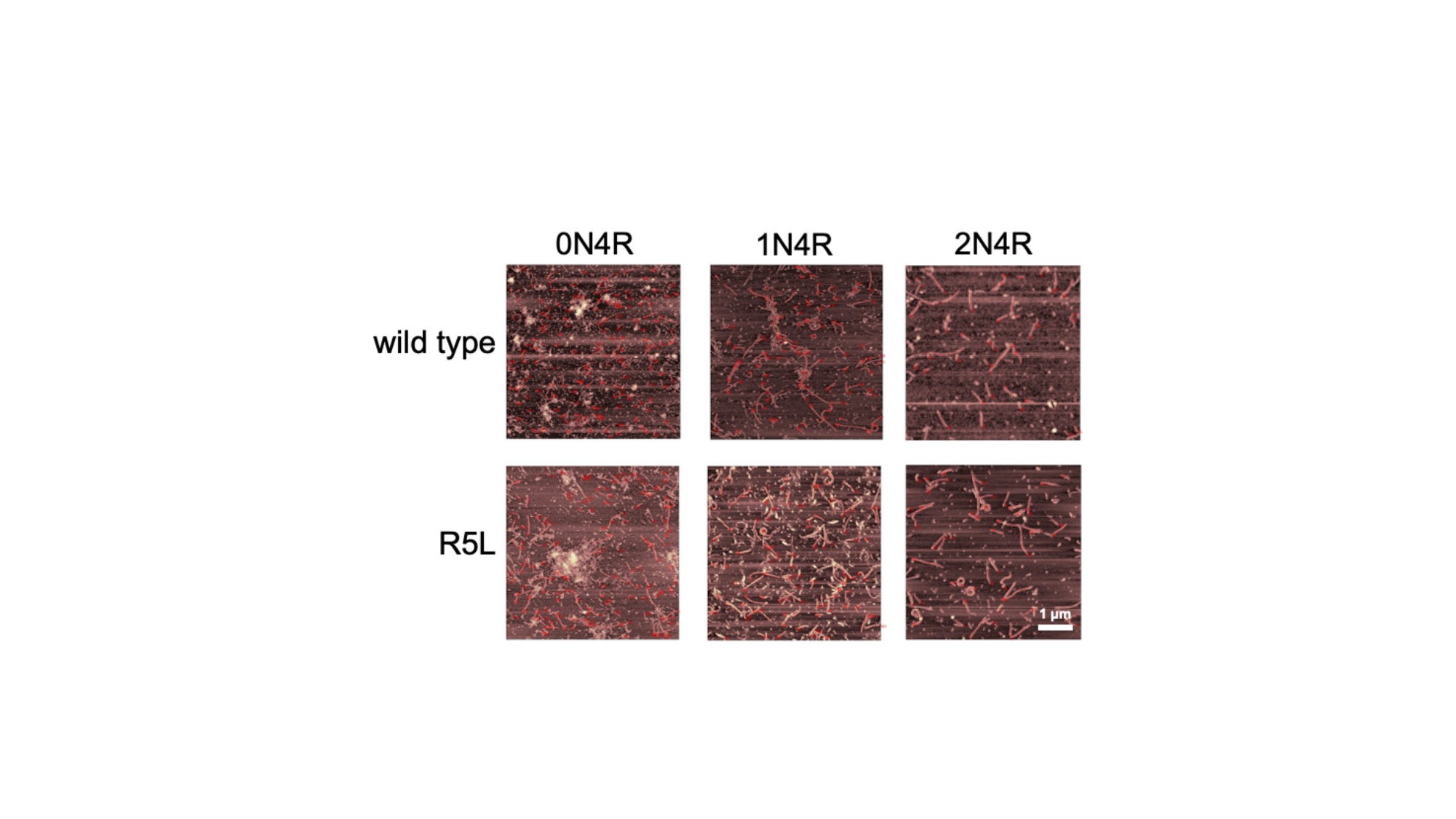

#### Slide 2
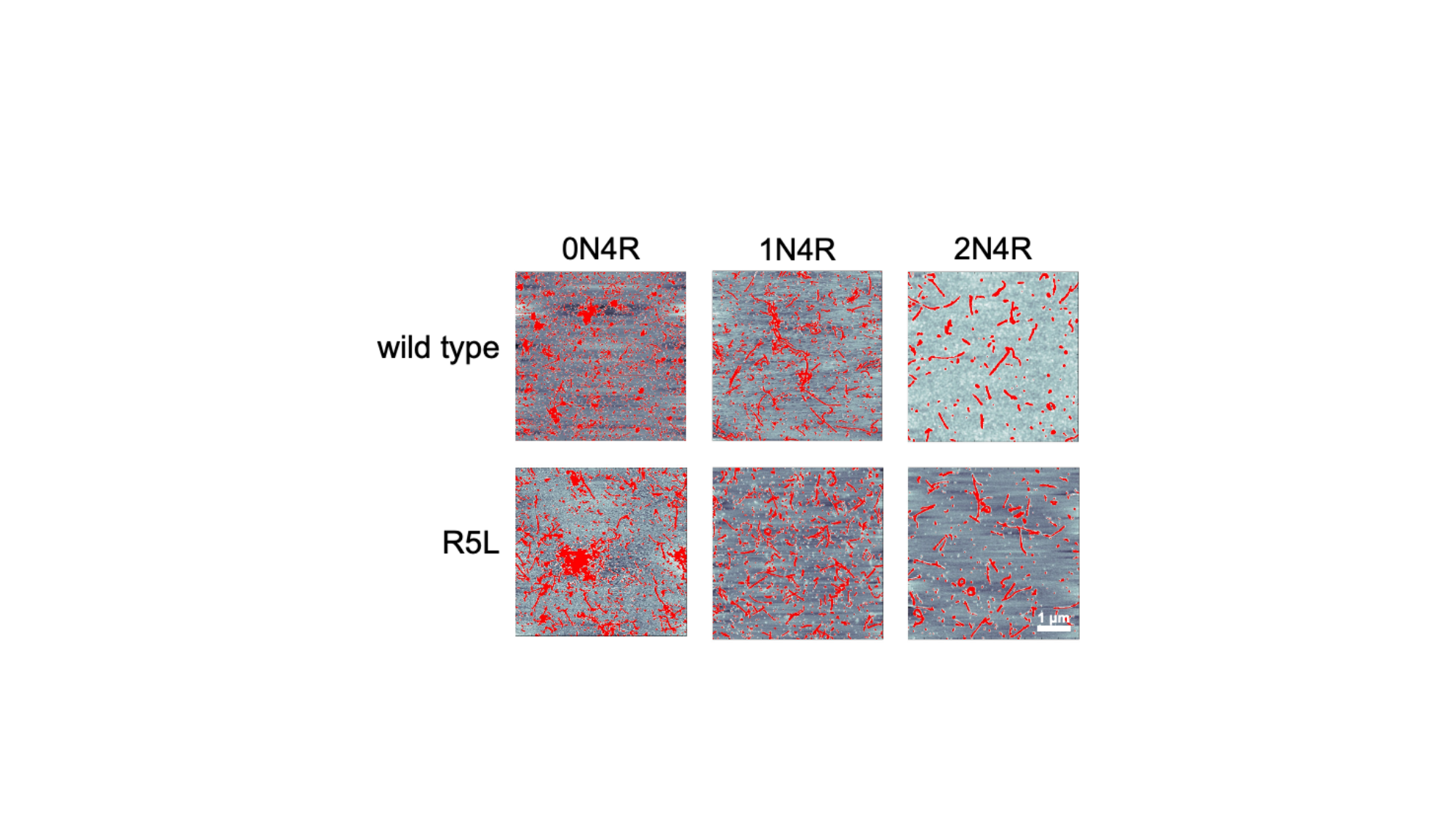

#### Slide 3
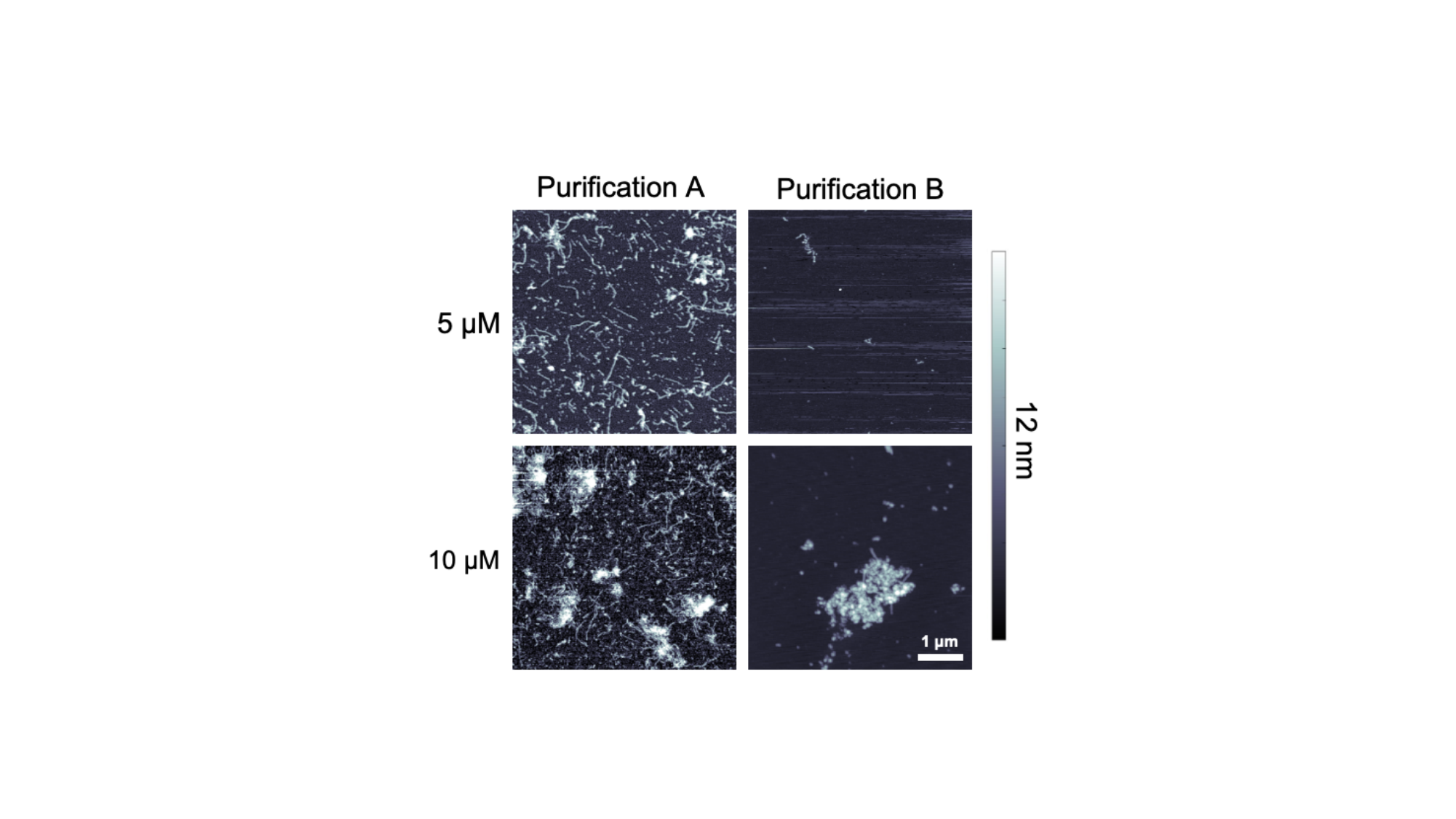

#### Slide 4
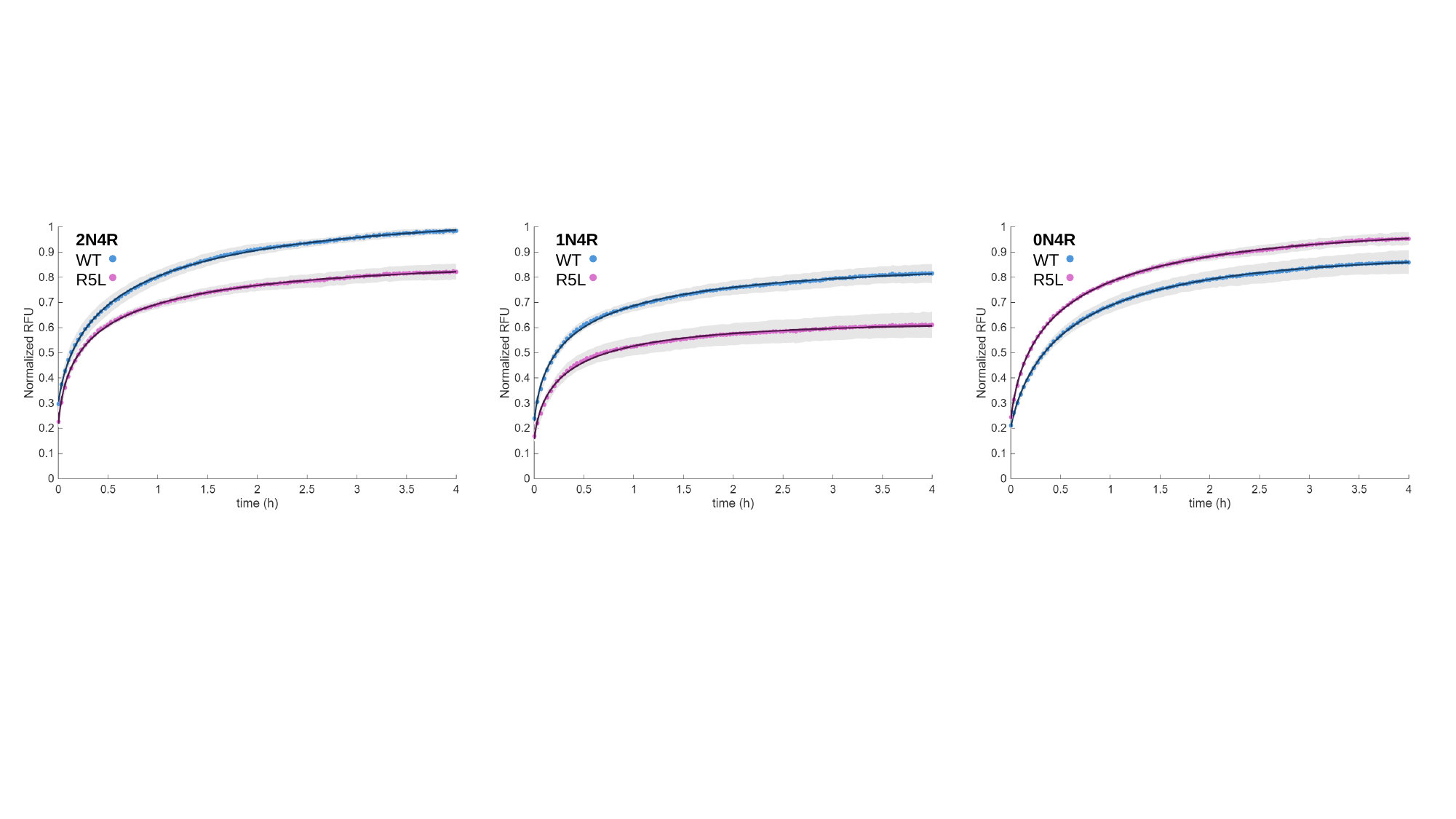

2N4R
WT
R5L
1N4R
WT
R5L
0N4R
WT
R5L

#### Slide 5
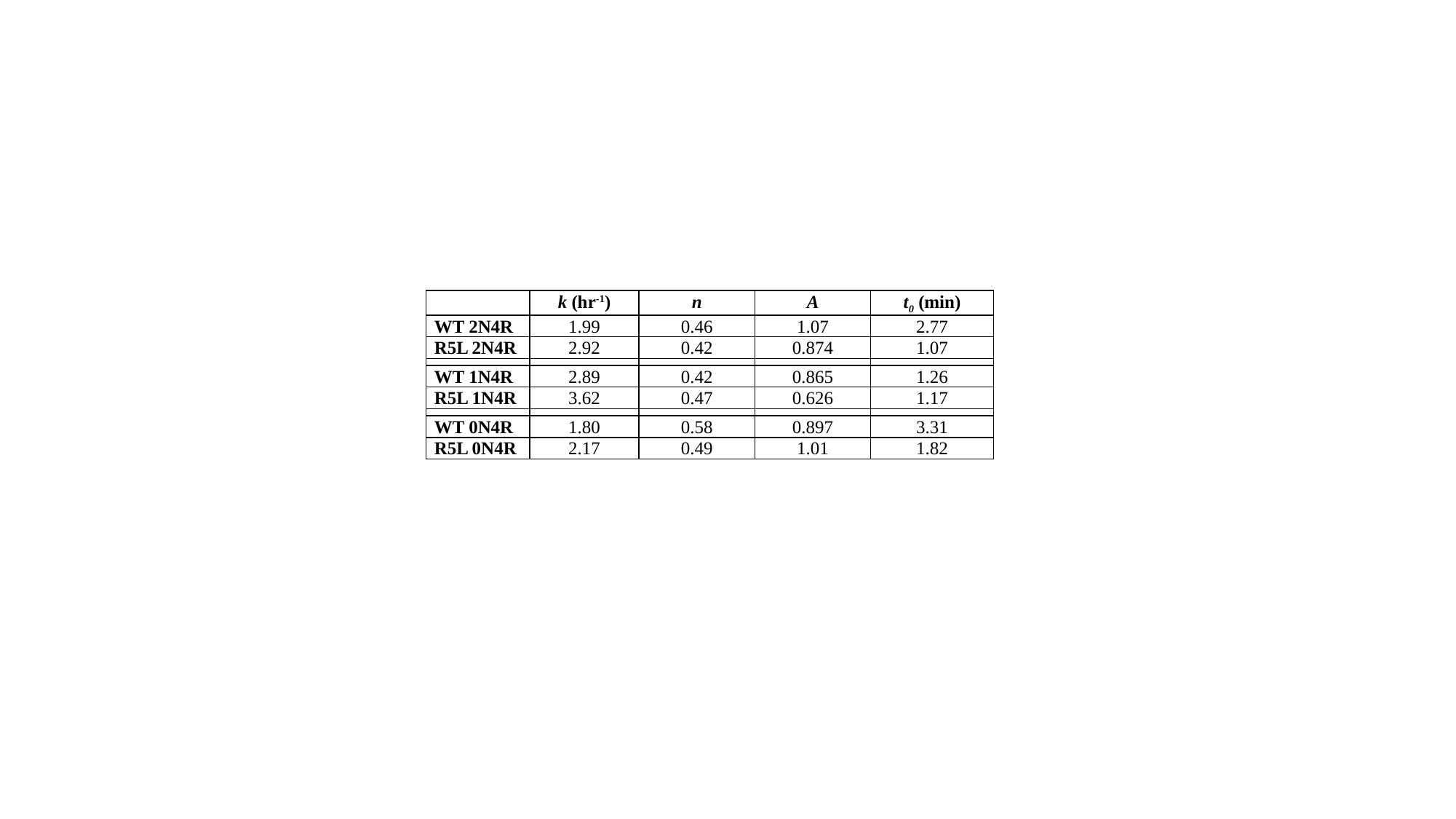

| | k (hr^-1^) | n | A | t_0_ (min) |
| --- | --- | --- | --- | --- |
| WT 2N4R | 1.99 | 0.46 | 1.07 | 2.77 |
| R5L 2N4R | 2.92 | 0.42 | 0.874 | 1.07 |
| WT 1N4R | 2.89 | 0.42 | 0.865 | 1.26 |
| R5L 1N4R | 3.62 | 0.47 | 0.626 | 1.17 |
| WT 0N4R | 1.80 | 0.58 | 0.897 | 3.31 |
| R5L 0N4R | 2.17 | 0.49 | 1.01 | 1.82 |
